# Antibiotics shift the temperature response curve of *Escherichia coli* growth

**DOI:** 10.1101/2020.04.04.025874

**Authors:** Mauricio Cruz-Loya, Elif Tekin, Tina Manzhu Kang, Alejandra Rodriguez-Verdugo, Van M. Savage, Pamela J. Yeh

## Abstract

Temperature variation—through time and across climatic gradients—affects individuals, populations, and communities. Yet how the thermal response of biological systems is altered by environmental stressors is poorly understood. Here we quantify two key features—optimal temperature and temperature breadth—to investigate how temperature responses vary in the presence of antibiotics. We use high-throughput screening to measure growth of *Escherichia coli* under single and pairwise combinations of 12 antibiotics across seven temperatures that range from 22°C to 46°C. We find that antibiotic stress often results in considerable changes in the optimal temperature for growth and a narrower temperature breadth. The direction of the optimal temperature shifts can be explained by the similarities between antibiotic-induced and temperature-induced damage to the physiology of the bacterium. We also find that the effects of pairs of stressors in the temperature response can often be explained by just one antibiotic out of the pair. Our study has implications for a general understanding of how ecological systems adapt and evolve to environmental changes.

## Introduction

Many environments experience daily and seasonal temperature fluctuations that affect rates of physiological processes. These changes in turn affect biological and ecological traits and ultimately impact the behavior of communities (1–9). In this manner temperature fluctuations can drive the evolution of organisms through variation in thermal sensitivity—the ability to function and survive at different temperatures (2, 10–15).

Measuring the growth of a living organism at different temperatures yields a temperature response curve (Figure 1a). Typically, temperature response curves have a single peak, corresponding to an optimal temperature where growth is maximized (2). As the temperature changes away from the optimum in either direction, the growth rate decreases, with an especially steep decline at higher temperatures. The range of temperatures in which an organism can grow to a certain extent (e.g. at least half of the maximum growth) is termed the temperature breadth. Living organisms are said to experience either cold or heat stress at extreme temperatures where growth is substantially less than optimum.

**Figure 1.**
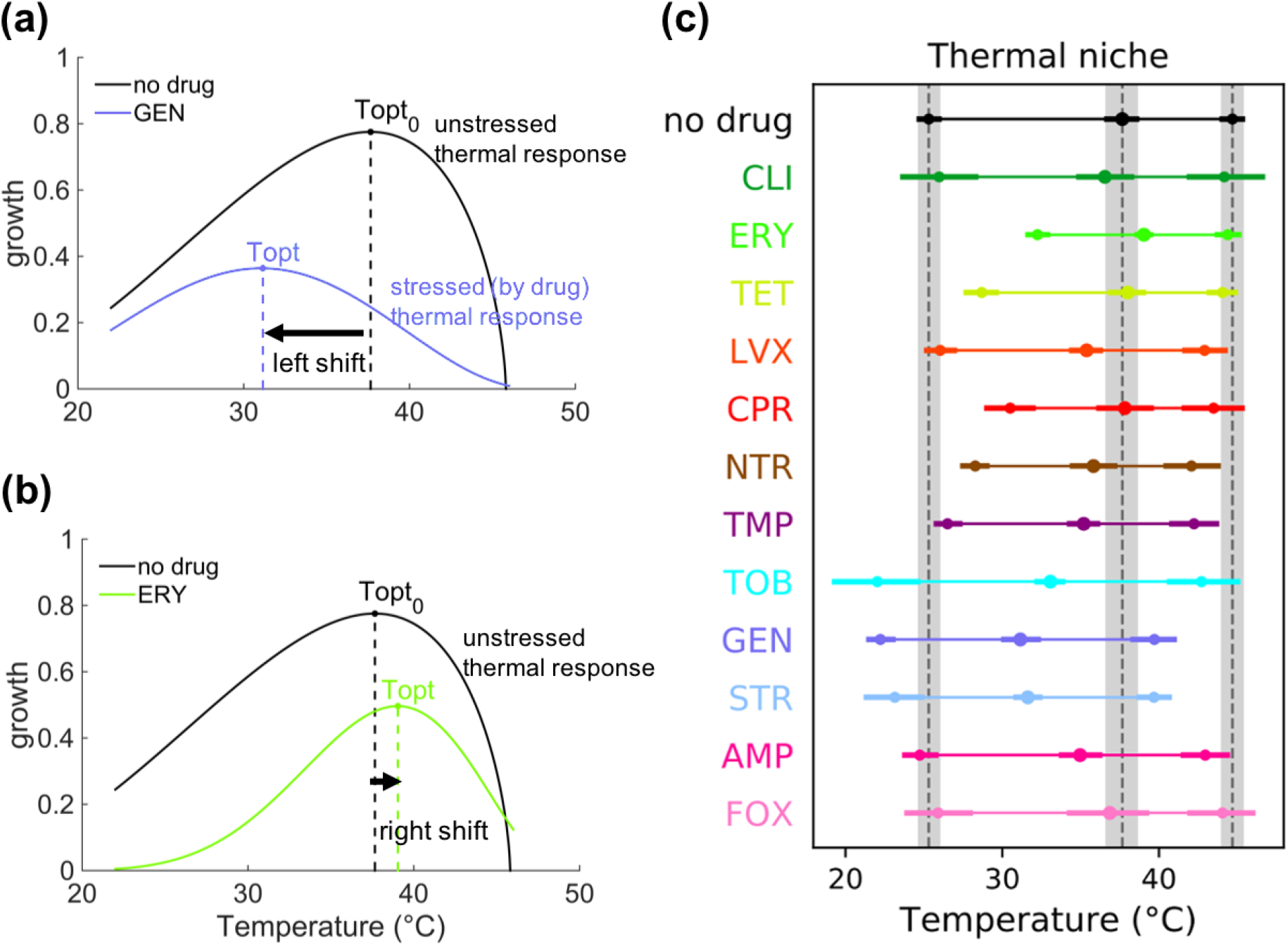
Temperature response curves change under antibiotic stress. **(a)** An example of a left shift of optimal temperature with antibiotic GEN. **(b)** An example of a right shift of optimal temperature with antibiotic ERY. **(c)** Optimal growth temperature (middle dot) and temperature niche (thin line joining the half-maximal growth temperatures, left and right dots) observed under each antibiotic used in this study. Point estimates for the optimal and half-maximal growth temperatures are shown as dots. To show the uncertainty in the estimates, 95% credible intervals (CIs, see Materials and Methods) are drawn as thick lines. The CIs for the no drug condition are shaded in the plot to facilitate comparison.

Thermal response curves are fundamental to grasping the variability of physiological and ecological traits in response to temperature changes in different taxonomic groups and habitats. Because shifts in the thermal response curves are representative of average fitness performance and temporal niches (16), optimal temperature and thermal breadth are indicative of evolution and acclimation patterns based on how species’ performance contributes to survivorship or fecundity (14). For instance, seasonal variation in temperature could lead to an evolution of different attack and escape speeds that would allow individuals to perform best when they are predator or prey (17).

At a cellular level the performance of an organism across different temperatures can lead to various genetic and physiological adaptation mechanisms. For example, in bacteria thermal sensitivity is related to many physiological and genetic modulations in metabolism, including outer membrane rigidity (18, 19), chemotaxis (20, 21), enzymatic thermo-stability (22, 23), and other general adaptive responses (24, 25). Moreover, the heat shock response—a cellular mechanism to deal with the deleterious effects of high temperatures, such as protein misfolding and aggregation—is highly conserved in both prokaryotes and eukaryotes (25, 26). Understanding responses to temperature changes is important to infer general patterns of how organisms, species, communities, and ecosystems are adapting to fluctuations in climate patterns and different environmental conditions.

Any environmental feature that kills a living organism or reduces its growth can be considered a stressor. Temperature can interact with other environmental stressors such as light, precipitation, pH, and salinity. Exposure to different stressor types and intensities can lead to phenotypic variation in an organism’s ability to respond to temperature changes (27, 28). Nevertheless, how the effects of environmental stressors interact with temperature responses is not well understood. Therefore, insights on whether temperature responses—as described by optimal temperatures and temperature breadths—can change rapidly and plastically in the presence of other environmental stressors have been lacking. In fact, it has been commonly assumed that thermal responses are not altered in the presence of other stressors (29–31).

A systematic approach that informs how optimal temperatures and temperature breadths are shifted by stressors (Figure 1) is needed to uncover these ambiguities and provide additional insights on fitness tradeoffs and thermal adaptation strategies. Here, we use a combined empirical-theoretical approach to study if the characteristics of thermal response curves change in response to additional environmental stressors. In particular, we use an experimental system of *Escherichia coli* and antibiotics as stressors in order to investigate how a physiological trait—growth of the bacterium—responds to variation in temperature in the presence of different stressor conditions. We obtain temperature response data for *E. coli* in 12 single-drug and 66 two-drug combination environments, where antibiotics are chosen to cover a wide range of mechanisms of action (Table 1). We then quantify both the optimal temperature and the temperature breadth of *E. coli* in the presence of these different environments.

**Table 1.**
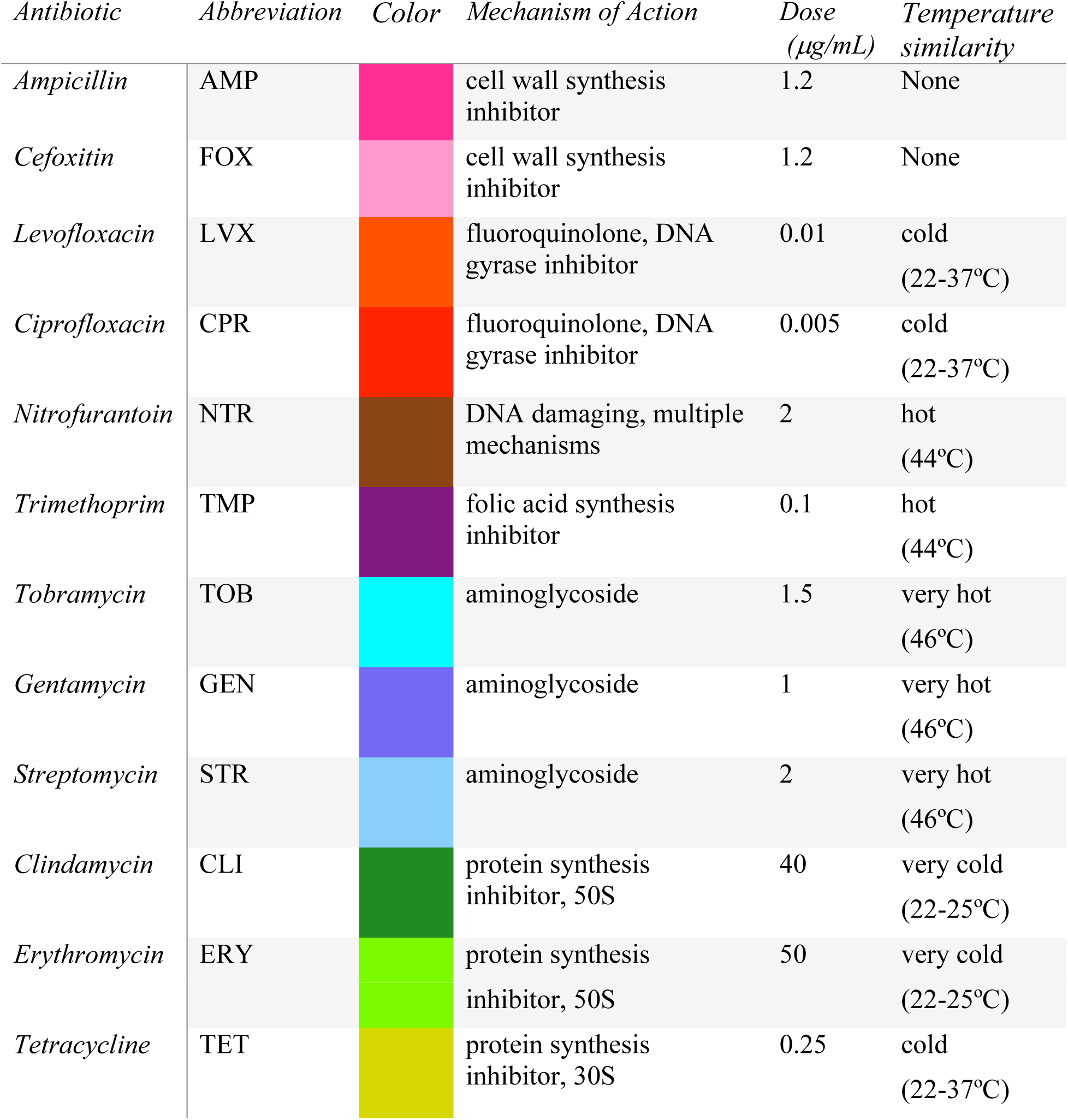
List of antibiotics. The antibiotics used are listed with their abbreviation, mechanism of action, dose, and our color scheme throughout the paper. Similar colors are chosen for drugs belonging to the same class/mechanism of action. For example, colors of blue tones are chosen for aminoglycosides. The similarity of each antibiotic to temperature stress according to their interactions with other stressors (32) is also shown, along with the corresponding range of growth temperatures that show similarity. For the purposes of cold/heat similarity, we consider any antibiotic with a similar temperature range lower than the optimum as cold-similar. For example, there are two groups of cold-similar antibiotics, which we call cold (22-37°C) and very cold (22-25°C). These terms are used to distinguish the groups by the relative strength of the cold stress they are similar to, but not necessarily the severity of the cold stress in an absolute sense.

We first show that individual stressors can have a substantial impact on the optimal temperature and temperature breadth. Next, we evaluate if the directions of the shifted thermal responses are related to the mechanism of action of the antibiotics. Previously, we determined that some specific classes of antibiotics have similar physiological effects to either heat or cold stress in *E. coli* (32). This classification was based on comparing the experimentally determined interaction profile (synergies and antagonisms with other stressors) of antibiotics and various growth temperatures (Table 1). We find that in most cases the direction of the shifts in the thermal responses under antibiotic stress can be explained through these groups. Lastly, we investigate how pairs of stressors move optimal temperatures in different directions as compared to the optimal temperatures under single-stressor conditions. In particular, we evaluate the extent to which the optimal temperatures result from integrated effects of both stressors, or whether a single stressor is the key driver of the temperature response. We infer from our results that a single drug often plays a dominant role in determining the optimal temperature response of a combined treatment.

Our experimental and theoretical framework on temperature response curves of *E. coli* presented here allows us to better understand how thermal sensitivities change in response to stressors. Therefore, our analysis will shed light on fundamental features shaping the ecological and evolutionary responses of organisms facing complex environmental conditions. By using antibiotics as stressors and a bacterium as a model organism, our study system is particularly valuable for its experimental tractability and reproducibility.

## Results

In this paper, we investigate how different stressors (antibiotics) alter an organism’s response to temperature, both in isolation and in combination. To do this, we determine the temperature optimum and temperature niche/breadth of *E. coli* by fitting the extended Briere model (developed here, see Materials and Methods) to experimental data of bacterial growth collected under different (unstressed and stressed) growth environments across multiple temperatures (22°C, 25°C, 30°C, 37°C, 41°C, 44°C, 46°C). The entire dataset and model fits are shown in Supplemental Figure 1. More details about the model and fitting procedure can be found in the Materials and Methods and Supplemental Information.

First, we explore how the optimal growth temperature of *E. coli* changes under single-stressor conditions (Figures 1a, 1b). We find that the majority of the single-drug environments exhibit left shifts—meaning the optimal temperature is lower—(Figure 1c) compared to the no-drug condition, *T*_*opt*_ = 37.7°C, CI: (36.7°C, 38.6°C). Right shifts are both less common and of lower magnitude than the observed left shifts. In addition, we find that the thermal niche breadth typically becomes narrower under antibiotic stress, meaning that *E. coli* can survive and properly function at a reduced temperature range.

Next, we investigate whether the physiological effects of antibiotics bear any relation to the direction of the observed shifts in the temperature responses (Figure 2). To do this, we group the antibiotics according to the similarity of their physiological effects to those of low or high temperatures, as determined previously (32). We observe the direction of the shifts for both single drugs (Figure 2a, left panels) and drug combinations that contain one or more of the antibiotics in the group (Figure 2a, right panel). We find that—for both single drugs and combinations—cold-similar antibiotics (i.e., with effects on bacteria similar to those caused by low temperatures) tend to either leave the optimal temperature unchanged or shift it slightly to the right (i.e., to higher optimal temperatures). In contrast, heat-similar antibiotics (i.e., with effects on bacteria similar to those caused by high temperatures) tend to result in unchanged optimal temperatures or shifts to the left (i.e., to lower optimal temperatures). In fact, bacteria exposed to aminoglycosides (TOB, GEN, STR), which induce misfolding of membrane proteins and have similar physiological effects to very high temperatures (Figure 2), show the greatest shifts towards the left. This is not the case for other protein synthesis inhibitors such as ERY or CLI that are similar to cold. Interestingly, beta-lactams shift the temperature curves in a similar way to heat-similar drugs when used in combinations, despite them having a different mechanism of action (inhibition of cell wall synthesis) that was not found to be heat-similar.

**Figure 2.**
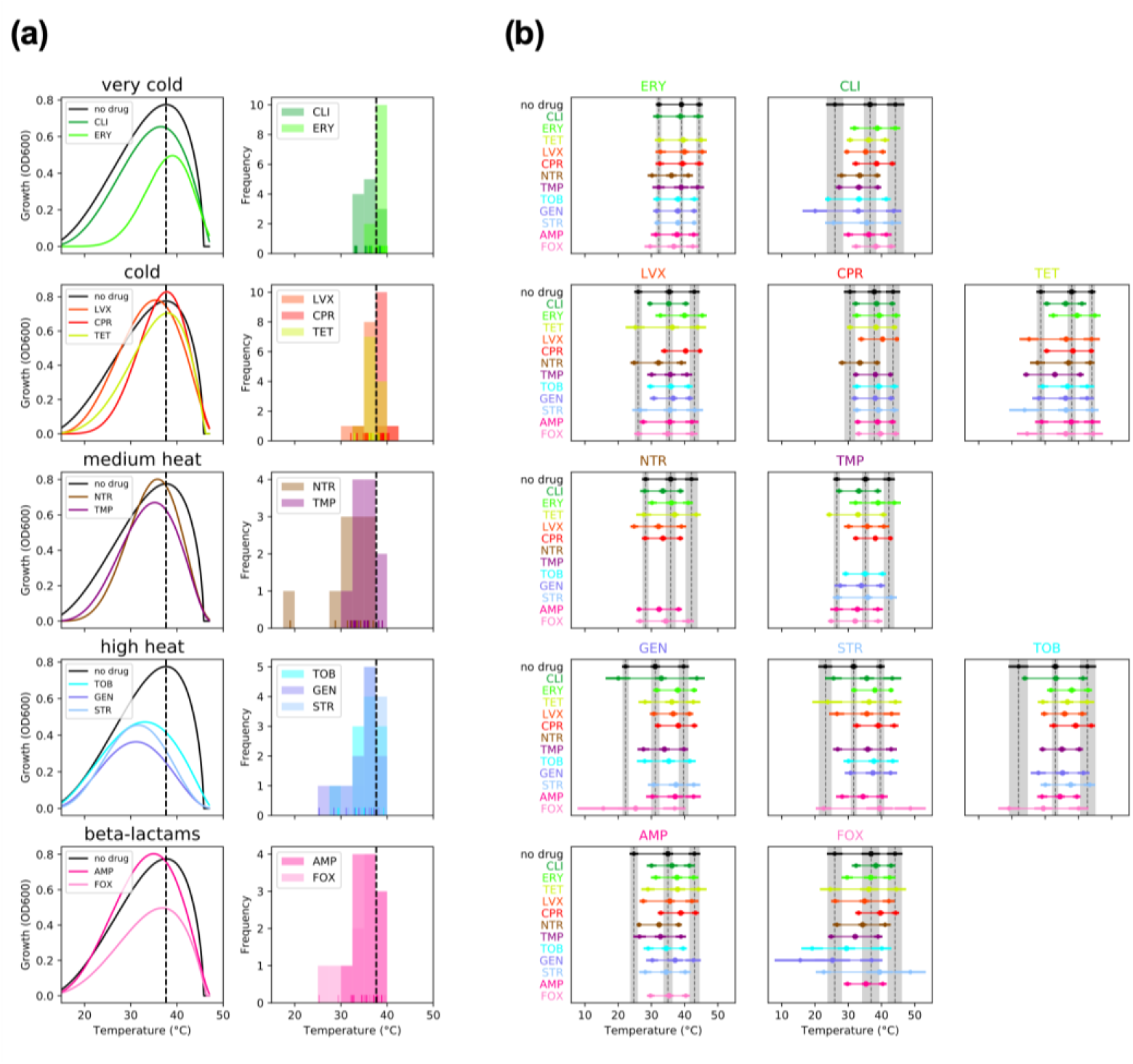
Physiological effects of antibiotics predict the direction of shifts in the optimal temperature. **(a) Left:** The fitted temperature response curve in the presence of single antibiotics is compared to the unstressed growth condition. Drugs are grouped according to the similarity of their effects to temperature (32), as shown in the top of the plots, except beta-lactams which did not show similarity to temperature. **Right:** Histogram of shifts in the optimal temperature under all pairwise drug combinations involving the drugs in the group. The individual estimates are shown as lines in the bottom. The unstressed optimal temperature is shown as a dotted line in both sets of plots. For both single drugs and combinations, the direction of the optimal temperature shifts depends on whether the drug is similar to cold or heat. **(b)** Optimal growth temperature and temperature niche observed under each antibiotic combination used in this study. The first drug in the combination is shown at the top of the plot. The second drug is shown in the y-axis using its assigned line color. The CIs for the single drug conditions are shown with shaded 95% credible intervals to facilitate comparisons. Conditions where the maximum growth was too small to estimate parameters reliably were removed.

We then compare the optimal temperature and temperature niche—the range between the temperatures that result in half-maximum growth—for bacteria under all antibiotic combinations to the single drug conditions (Figure 2b). For some antibiotics (e.g., ERY, CPR) the optimal temperature and the thermal niche range are similar to those of the single drug when combined with most other antibiotics. In contrast, there are other antibiotics for which these features show much more variation when combined with others (e.g. GEN, STR, TOB, FOX). This suggests that some antibiotics may act as the main drivers of the temperature response curve of antibiotic combinations.

Following this idea, we further explored how the optimal growth temperature is determined under combinations of stressors relative to the optimal temperature under single stressor conditions. We contrast the observed optimal temperatures with the predictions of five candidate models of how the combination optimal temperature could be determined from that of the single stressors (see Material and Methods, Figure 3a). The *min* and *max* models assume that the optimal temperature of the combination is determined by the optimal temperature of a single drug (the minimum or the maximum of the pair, respectively). These models best describe most (65%) multi-drug combinations (Figure 3b). The *attenuated* and *elevated* models assume that the optimal temperature of the combination is either lower or higher, respectively, than for both single drugs. These models best describe 18% of the combinations. Lastly, the *mean* model assumes that the temperature of the combination is determined by the average of the single drug optimal temperatures. This model best described only 17% of the drug combinations. These results suggest that the optimal temperature of antibiotic combinations is often determined by a single drug.

**Figure 3.**
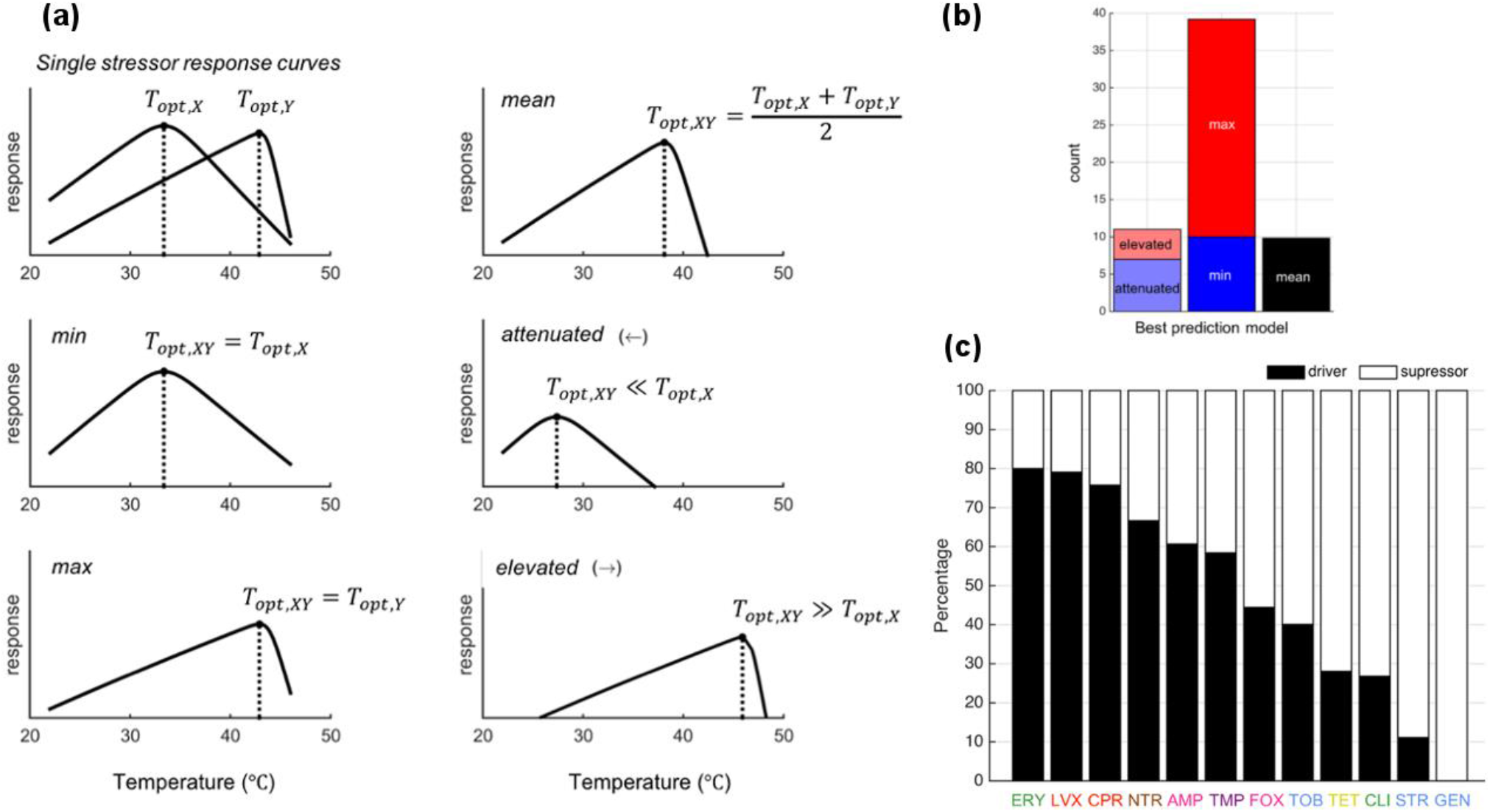
The optimal growth temperature under stressor combinations is often determined by a single stressor. **(a)** Schematic illustration of models to determine the optimal growth temperature under two stressors (*T*_*opt, XY*_) given the single stressor optimal temperatures (*T*_*opt, X*_, *T*_*opt, Y*_). **(b)** The frequency at which each model is the best fit, across all drug combinations. **(c)** Proportion of time each antibiotic is the main driver of the optimal temperature when combined with other antibiotics, based on the individual models: min, max, attenuated, and elevated.

Finally, we explore cases where single-driver models (*min*, *max*, *attenuated,* and *elevated*) represent the best optimal temperature model over the *mean* model, where both stressors compromise to result in the optimal temperature of an organism in the presence of combined stressors (Figure 3c). Interestingly, we rarely observe aminoglycosides (GEN, STR, TOB), antibiotics similar to high-heat, being drivers. In contrast, some cold-similar drugs (ERY, LVX, CPR), but not all (CLI, TET), frequently drive the optimal temperature of the combination. To account for the possibility that some antibiotics appear to be a driver more often than others purely by chance, we used a permutation test to evaluate our data against the null model that all drugs are equally likely to be a driver (see D-statistic in Supplemental Information for details). This test provides strong evidence (p=0.002) that some antibiotics have a greater tendency to be drivers than others by testing the entire dataset simultaneously. We also tested if specific antibiotics are drivers more often than expected by chance (see M-statistic in SI). However, we did not obtain statistically significant results for individual drugs, after correcting for multiple comparisons. We believe this may be due to a lack of statistical power to detect differences due to the smaller number of observations for individual antibiotics when compared to the full dataset.

## Discussion

Through a systematic analysis of growth response curves of bacteria across different temperatures and under different stressor environments, we investigate the effects of stressors on the phenotypic variation in temperature response traits—optimal temperature and temperature breadth. We see that stressors often decrease the temperature breadth and shift the optimal temperature in a direction that depends on their physiological mechanism of harm. In addition, our results suggest that left shifts—where the optimal temperature in a stressed environment is lower than the optimal temperature in unstressed environmental conditions—are more common and dramatic as opposed to right shifts towards higher optimal temperatures. This may be partially due to the asymmetry of temperature response curves, where the interval between the minimum and optimal growth temperatures is larger than that between the optimal and maximum growth temperatures.

High temperature harms living organisms through multiple mechanisms, including misfolding and aggregating proteins, damaging nucleic acids, and increasing membrane permeability (25). The heat-shock response attempts to prevent and/or repair this damage by producing chaperones that aid the correct folding of proteins (33). It has previously been shown that certain kinds of antibiotics can activate components of the heat-shock response (32, 34). However, adding antibiotics to heat stress is unlikely to help the cell survive high temperatures, since the heat-shock response is already induced by the high temperature alone. Thus, right shifts in the optimal temperature may be rare because it is unlikely that adding a second stressor can reduce or repair the high-temperature induced damage. In most cases where we do observe a right shift, it seems to be due to asymmetrical effects on the temperature response curve, where the left portion (i.e. below the unstressed optimal temperature *T*_*opt*_) is more depressed by the antibiotic than the right portion (Figure 2a, Supplemental Figure 1).

In contrast, cold temperatures predominantly slow down cell growth by suppressing DNA replication or protein translation (35, 36). Since the effects of low temperature seem to be primarily mediated by slowing down metabolism and growth rather than the accumulation of physical damage, it seems more likely that stressors can shift the optimal temperature to the left, especially when the stressor is more harmful at higher temperatures. In some cases, cold temperatures might allow cells to sustain antibiotic killing because certain antibiotics are only effective against actively growing cells (37). Low temperatures have also been shown to alter the structural stability (38) or the global uptake of some antibiotics such as gentamicin, thus impairing killing efficiency (39).

Based on network clustering methods (40, 41), we previously found that certain antibiotic classes have similar physiological effects to either heat or cold in *E. coli* (32). These temperature-drug groups were also shown to correlate with changes in drug sensitivity of high-temperature adapted strains obtained by Rodríguez-Verdugo *et al* (42). Interestingly, here we find that in most cases the direction of the shifts in the optimal temperature can be predicted from these groups. Cold-similar drugs tend to either leave the optimal temperature unchanged, or to shift it slightly to the right. In contrast, heat-similar drugs tend to result in larger shifts to the left or leave the optimal temperature unchanged (Figure 2). Similar trends are exhibited by antibiotic combinations containing drugs in these groups.

We propose a shared-damage hypothesis to explain this phenomenon: antibiotics that damage the same cellular functions as temperature stress (heat or cold) will cause an increased burden to the cell machinery that repairs this damage. For example, simultaneous exposure to aminoglycosides and high temperatures will result in more misfolded proteins than either stressor on its own.

Upon addition of an antibiotic, the stress-response machinery of the cell could be overwhelmed at less extreme temperatures, causing a greater reduction in growth at temperatures that cause similar physiological damage to the drug. The effect of these kinds of antibiotics in the temperature response curve will thus be asymmetrical. Growth will be more strongly reduced in the direction (heat or cold relative to Topt) where the drug and temperature damage overlap, and the optimal temperature will often shift towards the opposite direction because it suffers less from growth reduction. The cases with the most pronounced shifts in optimal temperature tend to have lower peak growth (Figure 2a). This suggests that perhaps these shifts become more pronounced when increasing the antibiotic concentration. Hence, we hypothesize that increasing the concentration of heat-similar drugs will result in greater shifts to the left and that doing so for cold-similar drugs will result in greater shifts to the right.

Notably, although the aminoglycosides (TOB, GEN, STR) share the same cellular target—the ribosome—as the other protein synthesis inhibitors (CLI, ERY, TET) used in our study, they result in distinct effects on the thermal response. Previously, differences in the effects of aminoglycosides and other protein synthesis inhibitors at different growth rates have been attributed to the reversibility of ribosomal binding (43). In that study, the authors found that STR is more effective when the growth rate is lower, which does not agree with our results at low temperatures. This discrepancy may be because the reduction in growth was previously manipulated by nutrient limitation as opposed to the temperature variation in our study.

Instead of binding reversibility, we could explain the different effects of these drugs by their mechanisms of action being qualitatively different, with the aminoglycosides being heat-similar and the other protein synthesis inhibitors being cold-similar. This is because aminoglycosides, unlike other protein synthesis inhibitors, induce mistranslation by the ribosome that decreases translational accuracy and causes protein misfolding (44). Cold temperatures may counteract this effect by slowing down ribosomal activity and increasing accuracy (45), thus causing aminoglycosides to be less effective when bacterial growth is suppressed at lower temperatures, as we observe. Reduced drug uptake at low temperatures could also play a role (39).

Interestingly, beta-lactams have a similar effect in the temperature response as heat-similar drugs. We speculate that this may be due to increased effectiveness at high temperatures due to a synergy between the cell-wall damage caused by the antibiotic and the increased membrane permeability caused by high temperatures. Further disentangling these processes in future studies will help increase our understanding of the connection between antibiotic susceptibility and bacterial physiology.

The breadth of the temperature niche is typically reduced in the presence of antibiotics, both in isolation and in combination. The lower and upper limits of growth are believed to be set by chemical and physical limits on the biological processes necessary for bacterial physiology, growth, and cell division (2). Since drugs introduce additional physiological damage in addition to that caused by temperature extremes, it seems likely that in most cases antibiotics would further constrain the temperature response and kill off barely-surviving populations at the extreme temperatures. Previous studies have also observed the temperature niche being reduced upon exposure to stressors (46, 47). From our results, it is apparent that stressors can reduce the temperature niche of living organisms at either temperature extreme. Thus, species that experience a wide range of stressful conditions at different times could perhaps experience selection for a broader temperature breadth that can be adapted to various environmental stressors (16).

The temperature niche is measured as the range between the low and high temperatures for half-maximum growth: its definition is therefore relative to the maximum growth. Consequently, conditions that decrease the right and central (i.e., near Topt) portions of the curve more than the left portion can result in a temperature niche that is shifted to the left (and vice versa) in the absence of increased growth at temperature extremes. These effects can lead to apparently surprising cases where adding a drug extends the limits for the thermal niche of the unstressed condition without increased growth at temperature extremes. This suggests that microbial communities may experience shifted thermal niches—giving rise to a different competitive landscape (e.g., due to reduced invasibility of high temperature habitats under aminoglycosides)—in the presence of certain antibiotics. These effects could be particularly important in the presence of variation in adaptations to antibiotics within microbial communities, which might cause the severity of their effects on the temperature curves to be species dependent.

We find that in most cases the shift in optimal temperature of *E. coli* due to antibiotic pairs is primarily determined by a single antibiotic. However, we also found cases where interactions between antibiotics seem to be important for determining the optimal temperature. For example, aminoglycosides (TOB, GEN, STR) show the largest degree of downshifting of the optimal temperature. However, when a second drug is added, this downshifting tends to be alleviated. Thus, in combinations of stressors, aminoglycosides are not the dominant driver for changing optimal temperature despite their large effects when used alone. When a shift in the thermal optimum is alleviated by addition of a second antibiotic, this does not imply that the reduced growth is also alleviated. Typically, these shifts are due to the second antibiotic decreasing some regions of the temperature response curve more sharply than others (Supplemental Figure 1). A notable exception involves interactions between certain aminoglycosides (GEN, STR) and other protein synthesis inhibitors (ERY, TET and other aminoglycosides), possibly because inhibition of protein synthesis reduces the aminoglycoside induced production of misfolded protein aggregates.

The growth response to multiple environmental factors such as temperature, CO_2_, and pH has been measured in green algae (48). In conditions with a large number of factors present simultaneously, the response is dominated by a single, severely detrimental driver. In contrast, in environments with a smaller number of factors, specific interactions between drivers were found to determine overall growth rather than the response to an overriding factor. These results were explained by the authors by the presence of a severely detrimental driver limiting the growth reduction that can be obtained by additional stressors, making the severe driver the primary determinant of the response. In contrast, we find that the effects of an antibiotic can sometimes be partially undone by another (e.g. aminoglycosides and other protein synthesis inhibitors). This suggests that, while identifying a dominant environmental driver can be a simplified approach to understanding organismal response to a complex system, this needs to be done with care since interactions between drivers can be a contributing factor as well.

The thermal optimum is often below the mean environmental temperature. This is because thermal response curves typically decrease sharply at high temperatures, so the penalty of going above the optimal temperature is much steeper than going below. Consequently, the exact distance between the thermal optimum and the mean is determined by the temperature variability in the environment (29–31). It is thus often tacitly assumed that the optimum temperature of individuals closely aligns the environment in which the individual has been reared and/or the species has evolved.

For this reason, the optimal temperature is not expected to quickly shift in response to other stressful conditions. This is a common assumption in mathematical models that describe the combined effects of temperature with other stressors such as pH (49), nutrient limitation (50), and humidity (51). In contrast, we observe that stressors can substantially and quickly change the optimal temperature for growth of a bacteria. A study that evaluated the combined effects of temperature and salt in slime molds (52) also found shifts in the thermal optimum, suggesting that this phenomenon is not limited solely to antibiotics.

Any physical or chemical environmental feature that kills or slows the population growth of a living organism can be considered a stressor. Antibiotics are stressors to bacteria in clinical settings but they may not always take this role in nature. It has been proposed that some antibiotics may participate in communication or be byproducts of metabolism in their natural environments (53–55). Since we explain thermal optimum shifts through differential growth reduction, our shared-damage hypothesis predicts antibiotic-induced thermal optimum shifts will occur when antibiotics are acting as stressors. However, this will not necessarily happen when antibiotics have a different role (e.g. communication) at much lower concentrations than those relevant in the clinic. In these cases, we would expect thermal optima to change only if there is a non-negligible fitness decrease caused by the antibiotic. Further work could test this by measuring the effects of an antibiotic on the thermal responses of microbes that naturally occur in the same environments as the antibiotic.

An exciting potential application of the shared-damage hypothesis is in predicting the effect of other stressors on the thermal optima of living organisms. To do this, further studies are necessary to evaluate the extent to which the physiological damage caused by other environmental stressors—such as pressure and pH—is similar to either temperature stress or antibiotics. This can be done by comparing either the gene-expression profile or the interaction profile (i.e. synergies and antagonisms with other stressors) of the environmental stressor of interest with those of extreme temperatures and/or antibiotics, as has been done to explore the overlap between antibiotics and temperature (32, 34). Our hypothesis would then predict that stressors that induce similar damage to high temperature will result in left shifts in *T*_*opt*_ (and vice versa). Moreover, the direction of the shift induced by a stressor should be the same as that of other stressors (e.g. antibiotics) that cause similar physiological damage. For example, beta-lactam antibiotics compromise the integrity of the bacterial cell wall so we speculate that the induced damage to the cell could have certain similarities to osmotic shock. If this were true, it seems possible that osmotic shock might change the temperature responses in a similar way to beta-lactams.

Although there has been substantial interest in understanding thermal response curves because of their potential to predict responses to climate change (17, 56, 57), the implications might be even broader. For example, an intriguing recent study showed that increased local temperatures were associated with increasing antibiotic resistance (58). This may be because temperature or seasonality effect environmental growth of resistant strains (59, 60) and horizontal gene transfer—one method of facilitating resistance transmission (61, 62). Another study showed that adaptations to long-term temperature changes unexpectedly coincided with mutations conferring resistance to rifampicin, an antibiotic that impairs RNA polymerase (42). Climate change has also been linked to changes on host-parasite dynamics that alter the frequency and severity of many infectious diseases (63, 64). Our work here and elsewhere shows that certain classes of antibiotics are more effective at different temperatures, and that there is substantial overlap in the response mechanisms to temperature and some kinds of antibiotics. This suggests the hypothesis that climate change might favor the evolution of resistance of specific (i.e. heat-similar) antibiotics indirectly by their resistance to high temperature stress.

From our results it also appears that drugs can be used to modify temperature response curves in predictable ways. A temperature-drug system could perhaps be used to examine scenarios for biological responses to climate change via a variety of thermal responses in a laboratory setting. Going forward, such a system could serve as a simplified model for examining changes in response to temperature across seasons, geographic gradients, and climate change.

Temperature is one of the fundamental drivers of biological processes. By using antibiotics as stressors, our study system is particularly valuable for its tractability, reproducibility, and potential to study temperature-stressor interactions beyond the pairwise level. Our results provide insights into the interactions between temperature and other stressors. Particularly, we show that stressors can modify the temperature response curves of a living organism, and that these changes can be predicted from the way the stressor harms its physiology. More broadly, our results imply that the chemical environment—or potentially the presence of other stressors—for a living organism can influence how it interacts with both abiotic (temperature) and biotic (modified competition due to changes in its thermal niche) factors. Investigating stressor effects on the physiological and ecological trait responses to temperature changes under this framework could lead to future research directions in exploring other environmental stressors that may aid in predicting the stability and diversity of ecological systems.

## Materials and Methods

### Experimental Framework

#### Bacterial strain and growth medium

The study used BW25113, a derivative of the F-, λ-, *E. coli* K-12 strain BD792 (CGSC6159) (65). Bacterial cultures were grown in LB broth (10 g/L tryptone, 5 g/L yeast extract, and 10 g/L NaCl) and maintained in 25% glycerol at −80°C. Fresh cultures were started by adding 20μL of thawed bacterial glycerol stock into 2 mL of LB followed by incubation at 37°C. Cultures were grown to exponential growth phase and diluted to maintain 10^4^ cells per experimental condition.

#### Antibiotics

A total of 12 antibiotics were included in the study as representatives of all major drug classes. Ciprofloxacin (CPR) from MP Biomedicals (Santa Ana, CA) and Gentamycin (GEN), levofloxacin (LVX), tetracycline (TET), tobramycin (TOB), erythromycin (ERY), ampicillin (AMP), clindamycin (CLI), streptomycin (STR), nitrofurantoin (NTR), cefoxitin (FOX), and trimethoprim (TMP)—all from Sigma (St Louis, MO)—were used. Stock solution at 20 mg/mL of each antibiotic was stored in 50μL aliquots at −20°C. Each aliquot was only frozen and thawed once to preserve potency.

#### Growth experiments

Antibiotics used in all experiments inhibited bacterial growth at sub-lethal concentrations (50% to 90% growth). The desired concentrations were first determined by a twelve-step concentration series of 2-fold at each step in 96-well plates (Costar). Antibiotic stock solutions were prepared in a total volume of 5 mL at 10-fold of their respective concentrations (Table 1). Experiments of pairwise drug combinations were prepared by adding 10μL of each component drug followed by the addition of 80μL cell inoculum. 10μL of LB medium was added in replacement of a second drug for single drug experiments. Each experimental condition was conducted in 4 replicates from the same antibiotic stock solution. The plates were incubated at various temperatures (22°C, 25°C, 30°C, 37°C, 41°C, 44°C, 46°C) with aeration at 300 rpm. Cell density was measured at 4-hours, 8-hours, 12-hours and 24-hours by reading at OD600 nm. The optical density measurements (used as a proxy for bacterial growth) at 24-hours were used to infer the temperature curves.

### Mathematical Framework

#### Extended Briere model for characterizing temperature response curves

Briere (66) defines a simple model for the temperature dependence of a trait, such as growth, denoted by *g*(*T*) as follows:

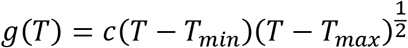

where *T*_*min*_, *T*_*max*_ are the minimum and maximum temperature of growth. This equation can be solved analytically (see Supplementary Material) to show that the optimal temperature yielding maximum growth is always attained at 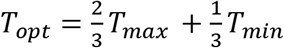. This model is not flexible enough to describe the antibiotic growth curves we found empirically. As a more general alternative, we propose an extended Briere model

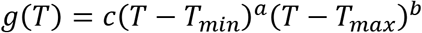

where *a*, *b* ≥ 0 are parameters that determine the shape of the curve. In this extended Briere model of temperature dependence of a trait, we have *T*_*opt*_ = *αT*_*max*_ + (1 − *α*)*T*_*min*_. In this model the optimum temperature can lie anywhere between *T*_*min*_ and *T*_*max*_ depending on the value of the fraction 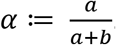.

The extended Briere model can be reparametrized as

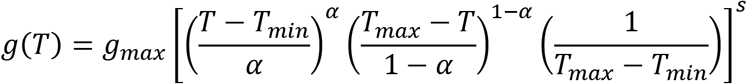

where *g*_*max*_ is the maximum value of the trait, i.e. growth, and *s* = *a* + *b* is a parameter that determines the smoothness of the temperature response curve (see Supplementary Material for details). We use this parametrization for parameter fitting of temperature response curves of the bacterium across different drug combination treatments (see Table 1 for chosen drugs in our study).

#### Bayesian parameter fitting

The extended Briere model was fitted to the temperature growth curve for the bacterium under all conditions through a Bayesian methodology with the `pymc3` library of the Python programming language (67). The following methodology is used for obtaining Bayesian estimates for the model parameters. Let *y*_*i*_ be the *i*th observed data point for growth after 24 hours and let *T*_*i*_ be the temperature at which it was observed. The observed values were assumed to be Gamma distributed with

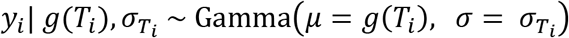

where the Gamma distribution is parametrized in terms of the mean *μ* and standard deviation *σ*. The data are clearly heteroskedastic, and multiple measurements were taken at the same temperature enabling estimates of the standard deviation at each measured temperature. Because of this, a different standard deviation was modeled for each measured temperature. The following hierarchical model was used for the standard deviation:

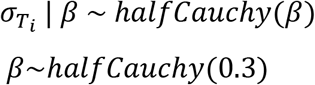

The prior distributions for the extended Briere model parameters and a justification for all priors used is given in the Supplementary Material. A variational method (full-rank ADVI) (68) was used to obtain approximate posterior distributions for the model parameters. These posterior distributions were used to construct point estimates—the expected value of the posterior distribution—and 95% credible intervals for all parameters to evaluate the uncertainty in the estimates. Credible intervals are the Bayesian analog to confidence intervals. A credible interval contains the true value of the parameter of interest with the specified (e.g. 95%) probability, given the observed data.

#### Models for predicting optimal temperatures for multi-drug responses

We denote single drugs as *X* and *Y*, and the combination of drugs as *XY.* We use these drug notations as a subscript for the corresponding optimal temperatures, i.e. *T*_*opt,X*_, *T*_*opt,Y*_, and *T*_*opt,XY*_. For predicting the optimal temperature of multi-drug combination treatments, we define five different models by the choice of simple yet biologically meaningful scenarios (Figure 3).

i. A single drug is playing a major role in determining the optimal temperature of the bacterial response. The optimal temperature of the combination is given by either of the two individual stressors (*min* or *max* model). *T*_*opt,XY*_ = min(*T*_*opt,X*_, *T*_*opt,Y*_) or *T*_*opt,XY*_ = max(*T*_*opt,X*_, *T*_*opt,Y*_).
ii. The optimal temperature of the bacterium in the presence of drug combinations is shifted to lower or higher temperatures than both single drugs’ optimal temperature values. Along the lines of these extreme behaviors, we define *attenuated* and *elevated* optimal temperature models *T*_*opt,XY*_ ≪ min(*T*_*opt,X*_, *T*_*opt,Y*_) or *T*_*opt,XY*_ ≫ min(*T*_*opt,X*_, *T*_*opt,Y*_).
iii. Temperature tolerance is determined by both of the drugs in the combination. To uncover such cases, we define our fifth model, namely the *mean* optimal temperature model. This model expresses the optimal temperature of the combined treatment as the average of the two single drug optimal temperatures (Fig 1). In other words, the *mean* model is equivalent to 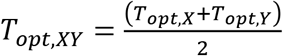.

To determine the best optimal temperature model, we measured the difference between the actual value and the predicted value of each of the *min*, *mean,* and *max* models. We considered the best-fit model as the one with the smallest absolute difference between actual and predicted values. When this absolute difference is greater than the cutoff value of 2.20°C (see Supplementary Figure 2), then the best model is determined to be either the *attenuated* or *elevated* model depending on the direction of the optimal temperature shift.

## Data availability

The dataset and code used in this manuscript will be made available upon publication and for peer review upon request.

## Acknowledgements

We thank Nina Singh for comments on the manuscript. We thank Rina Watanabe for laboratory assistance. We are grateful for funding from the Hellman Foundation (PJY), a KL2 Fellowship (PJY) through the NIH/National Center for Advancing Translational Science (NCATS) UCLA CTSI Grant Number UL1TR001881, UC Mexus and CONACYT (MCL), and a James F. McDonnell Foundation Complex Systems Scholar Award (VMS).

---

**Supplemental Figure 1.**
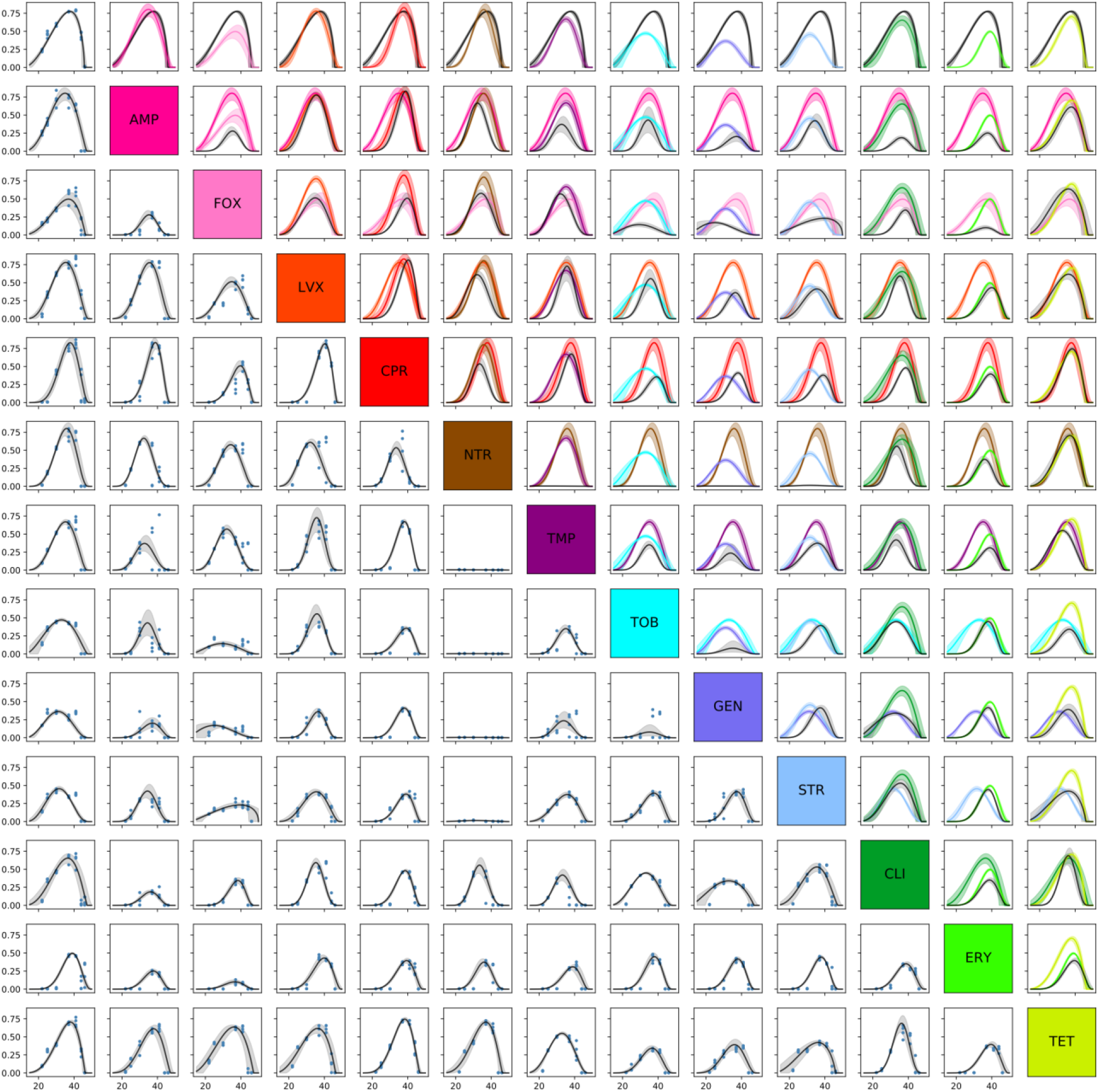
Full dataset and model fits. **Lower half:** The growth data for all antibiotic combinations (blue dots), as well as the fitted extended Briere model (black lines), are shown, as well as 95% credible intervals for the extended Briere model. The upper left corner corresponds to the no drug case. **Upper half:** The fitted curves corresponding to each drug combination are shown in black, and the fits corresponding to each single drug are shown in their corresponding color. In the uppermost row, the growth curve in the absence of antibiotics is shown in black and the single drug curves are shown in the corresponding color.

**Supplementary Figure 2.**
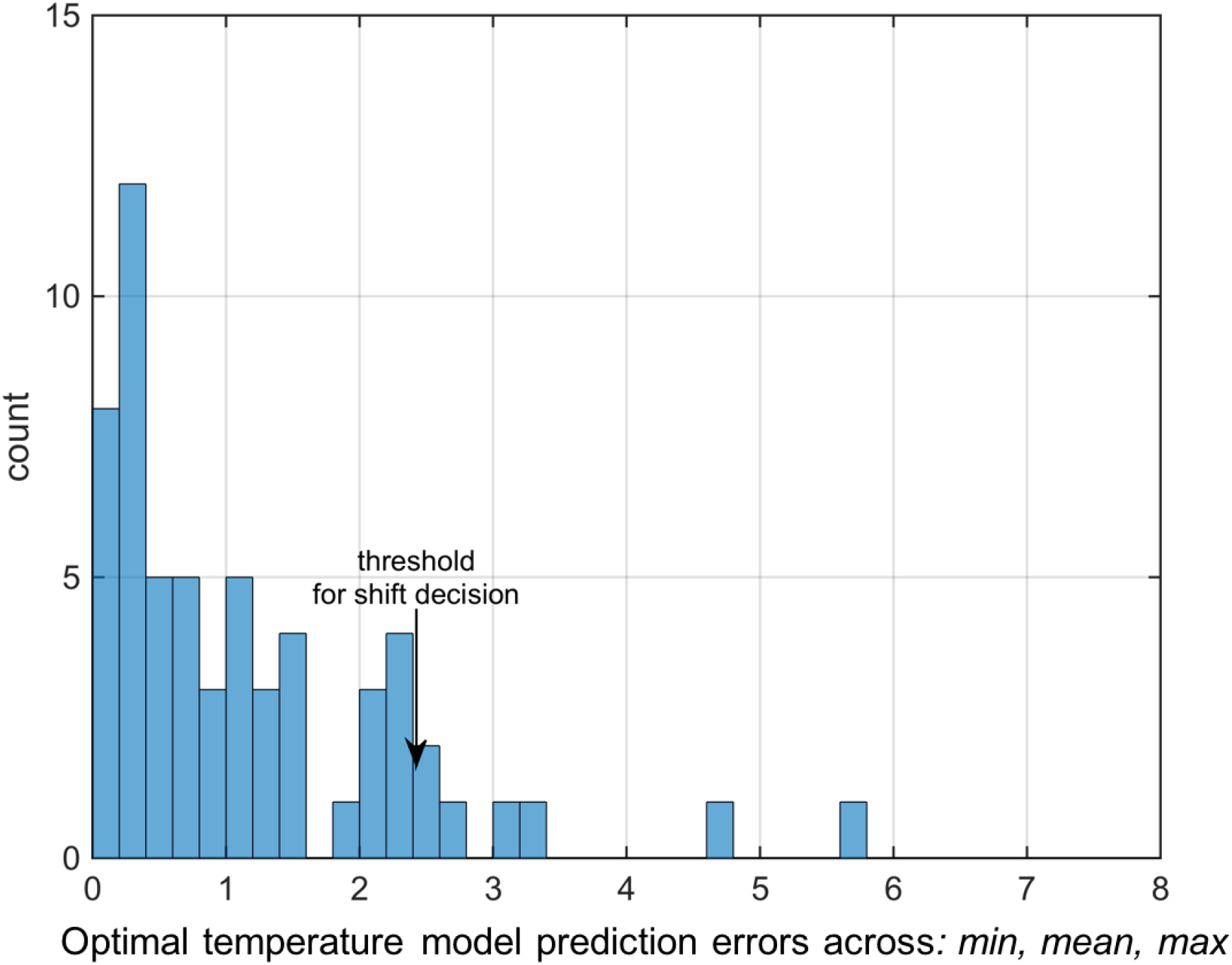
Threshold for distinguishing single-driver models (*min* versus *attenuated* or *max* versus *elevated*). The distribution of model prediction errors across the *min*, *mean*, and *max* models is plotted to decide a cutoff to determine if the optimal temperature shift of a pairwise combination is large enough to decide the attenuated or elevated models are a better fit. When the prediction error (as defined by an absolute value of difference of optimal temperature prediction and actual optimal temperature) is higher than 2.20 °C, the best model is either the attenuated model or elevated model based on the direction.

**Supplementary Figure 3.**
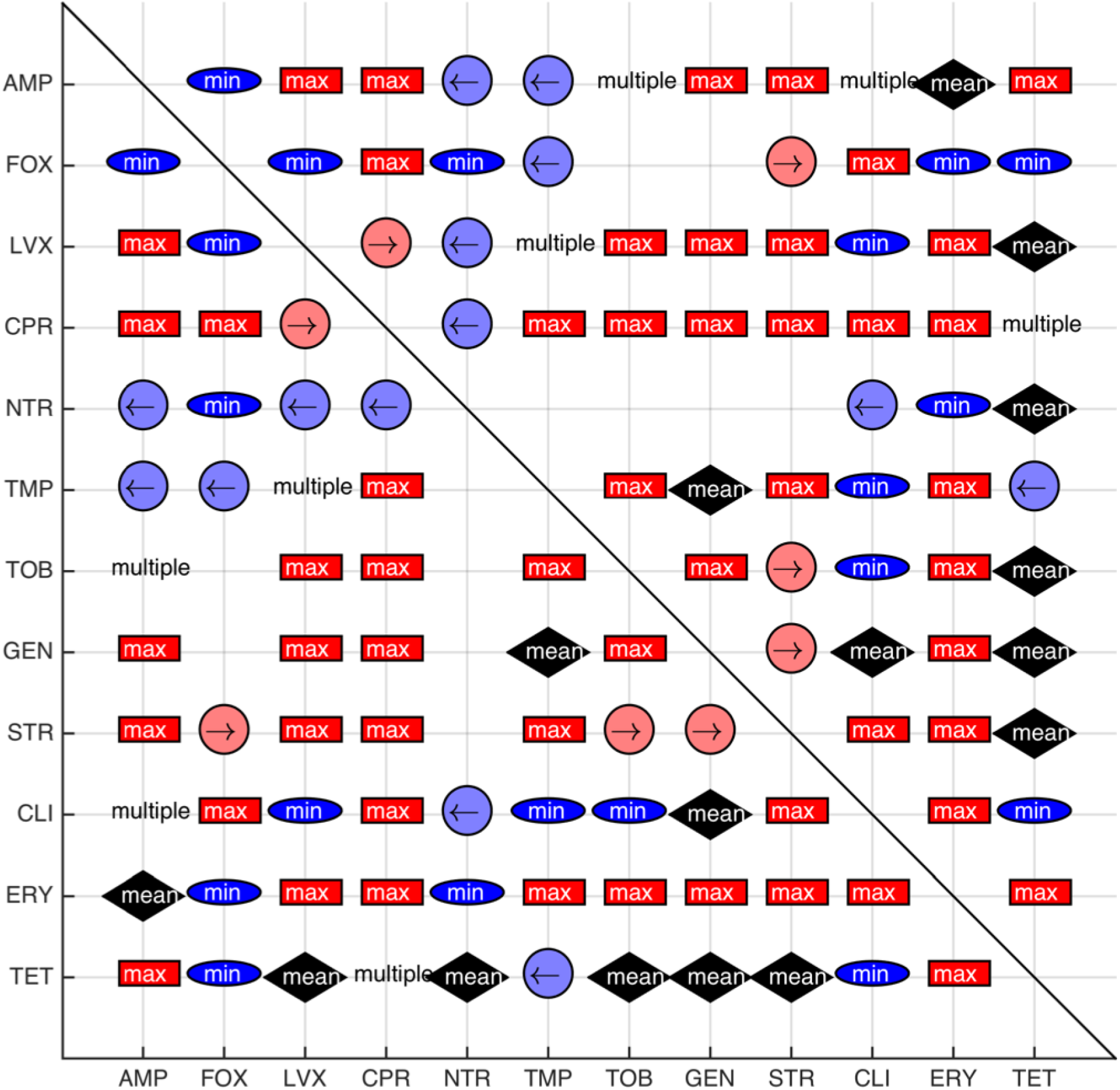
The optimal temperature model yielding the best prediction for each drug pair is shown. The blue oval frame represents the *min* model, the red rectangular frame represents the *max* model, the light red circle frame with the arrow pointing right represents the *elevated* model, and the light blue frame with the arrow pointing left represents the *attenuated* model. Four pairs—specifically AMP+TOB, CLI+AMP, TMP+LVX, and CPR+TET—exhibited multiple models showing the best model prediction, with all results showing *mean* and *max* to be the best models. These pairs are displayed as “multiple” in the figure.

